# Patterns of Ongoing Thought in the Real World and Their Links to Mental Health and Well-Being

**DOI:** 10.1101/2024.07.22.604681

**Authors:** Bridget Mulholland, Louis Chitiz, Raven Wallace, Brontë Mckeown, Michael Milham, Arno Klein, Robert Leech, Elizabeth Jefferies, Giulia Poerio, Jeffery Wammes, Jeremy Stewart, Samyogita Hardikar, Jonathan Smallwood

## Abstract

The thoughts we experience in daily life have implications for our mental health and well-being. However, it is often difficult to measure thought patterns outside of laboratory conditions due to concerns about the voracity of measurements taken in daily life. To address this gap in the literature, our study set out to measure patterns of thought as they occur in daily life and assess the robustness of these measures and their associations with trait measurements of mental health and well-being. A sample of undergraduate participants completed multi-dimensional experience sampling surveys eight times per day for five days as they went around their normal lives. Principal component analysis reduced these data to identify the dimensions that explained the patterns of thought reported by our participants. We used linear modelling to map how these thought patterns related to the activities taking place at the time of the probe, highlighting good consistency within the sample, as well as substantial overlap with prior work. Multiple regression was used to examine associations between patterns of ongoing thought and aspects of mental health and well-being, highlighting a pattern of “Intrusive Distraction” that had a positive association with anxiety, and a negative association with social well-being. Notably, this thought pattern tended to be most prevalent in solo activities and was relatively suppressed when interacting with other people (either in person or virtually). Our study, therefore, highlights the use of multi-dimensional experience sampling as a tool to understand how ongoing thoughts in daily life impact on our mental health and well-being, and establishes the important role social connectedness plays in the etiology of intrusive thinking.

## Introduction

A fundamental goal of psychology is to understand how the thoughts we experience in daily life relate to our productivity, mental health, and well-being. Contemporary work indicates a link between patterns of ongoing thought, as determined by experience sampling, and various aspects of health and well-being (e.g., [1, 2]). However, research on human thought has historically relied on observations made in controlled laboratory contexts such as brain imaging (e.g., [3]) or behavioral laboratories (e.g., [4]) because of the assumption that the robustness of these measures is improved when they are gathered in controlled conditions. However, recent work in the laboratory and in daily life has emphasized that patterns of thought are intimately related to the context in which they emerge [5]. Since the tasks people perform in the lab are unlikely to perfectly match the activities people perform in daily life, the role that context plays in influencing thought patterns may be one reason why links between traits and patterns often do not generalize from the laboratory to daily life [6].

The influence that context has on thought patterns has been established both within the laboratory and in daily life (for a review see [7]). For example, Konu and colleagues [8] used a battery of laboratory-based activities, including cognitive and attentional tasks, videos, and audiobooks to establish that the patterns of thought people engaged in changed as a function of the tasks people engage in. In a similar vein, Mulholland and colleagues [9] established that in daily life, the thought patterns participants reported varied with the activities people engaged in. There were also similarities in how the ongoing context influenced the patterns that participants reported. In both studies, patterns of thought with (i) unpleasant, intrusive, and distracting features were identified that dominated low-demanding situations in both the lab and daily life, (ii) thoughts with social and episodic features were important in social situations in daily life and tasks relying on social cognition in the lab, and (iii) patterns of detailed focus on a task were important during homework and while working in daily life, and were present during demanding laboratory task such as those depending on working memory. These prior lab and daily life studies [8, 9] took advantage of a novel approach to measuring cognition known as multi-dimensional experience sampling (mDES, [10]). This approach asks participants to rate their experience along a number of dimensions (e.g. how detailed they are, whether are they intrusive, whether they involve other people, etc.…) on multiple occasions. These data are then often decomposed into a set of underlying dimensions that describe the patterns of answers that the participants provided. This dimension derived from this decomposition provides a ‘thought space’ in which different experiential moments can be located [11–13]).

Emerging evidence suggests that one reason why mDES can characterize contextual influences on ongoing thought is because it provides high precision descriptions of individuals’ experiences. For example, states identified using mDES are associated with brain activity during attention tasks, where patterns off-task thought with episodic features are related to greater activity within the medial prefrontal cortex, a region of the default mode network [14]. In contrast, patterns of detailed focus relating to a task are linked to greater activity in the frontoparietal network [12]. mDES is also sensitive to changes in brain activity emerging during movie watching [11], highlighting that patterns of intrusive distraction are related to moments during a film when frontoparietal activity is reduced [11], a pattern consistent with the notion that intrusive thought reflects a failure to control cognition [15]. Recently, we demonstrated that mDES is sufficiently sensitive to variation in thought patterns to the extent that it can be used to build a fully generative model of the mapping between thought patterns and brain activity during task states [13].

### Current Study

Together, these prior studies demonstrate that mDES provides a powerful tool for mapping thinking patterns in the lab and daily life. Thus, our goal was to understand how the thought patterns mDES reveals in daily life relate to measures of mental health and well-being. Prior studies have already used experience sampling in daily life and in the laboratory to highlight important links between patterns of ongoing thought and factors that impact mental health and well-being, such as depression [16, 17], anxiety [18], and obsessional thinking [19]. The goal of the current study was to leverage the precision mapping of experience sampling offered by mDES to understand the links between thought patterns in daily life and mental health and well-being. However, before examining links to mental health and well-being, we first investigated the reliability of the thought patterns produced using mDES. We took advantage of the fact that the thought patterns established by mDES are sensitive to the activities being performed in daily life [9] and examined whether the mappings between thought patterns and activities are consistent within the current dataset and also with those seen in our prior study [9]. Once we established the reliability of mDES as a tool for mapping thought patterns in daily life, we examined how the thought patterns established by our approach related to aspects of mental health and well-being as measured by a battery of questionnaires.

## Methods

### Participant Population

A total of 261 participants (women = 227, men = 29, non-binary or similar gender identity = 4, prefer not to say = 1; *M* = 21.44; *SD* = 6.16; range = 17-52; note, two ages were recorded incorrectly and therefore not included and two additional ages were absent) were included in this study. This study was granted ethics clearance by the Queen’s University General Research Ethics Board. Participants were recruited between January 9th, 2023, and April 10^th^, 2023, through the Queen’s University Psychology Participant Pool. Eligible participants were Queen’s University students enrolled in designated first- and second-year psychology courses. Participants gave informed, written consent through electronic documentation prior to taking part in any research activities and were awarded two course credits and fully debriefed upon the completion of their participation.

### Procedure

Participants completed demographic questionnaires (i.e., age, gender, gender identity, sex assigned at birth, language(s) spoken most often at home, country of primary place of residence, program, and year of study). Participants also completed evidence-based health and wellness questionnaires, including the Quick Inventory of Depressive Symptomatology self-report (QIDS-SR_16_) [20], the Anxiety Sensitivity Index (ASI) [21], the Autism Spectrum Quotient (AQ) [22] to assess for autistic traits, the Adult ADHD Self-Report Scale (ASRS) [23] to asses for ADHD traits, and the Montgomery Asberg Depression Rating Scale self-report (MADRS-S) [24] to measure severity of depression symptomatology and the Overall Anxiety Severity and Impairment Scale (OASIS) [25] to measure severity of anxiety symptomatology, the MOS 36-item short form survey (SF-36) [26], the Sheehan Disability Scale (SDS) [27] to assess functional impairment, and the World Health Organization Quality-of-Life Brief Version (WHOQOL-BREF) [28] to quantify quality-of-life. These questionnaires were selected based on a widespread, general approach to health and wellness, rather than clinical diagnosis. For the purpose of this study, our analysis was limited to the AQ, ASRS, MADRS, OASIS, and WHOQOL-BREF questionnaires which were deemed the most relevant indicators of mental health and well-being. Once participants completed these questionnaires, they were notified via Samply Research, a mobile application, to complete mDES questionnaires eight times daily for five consecutive days between 7:00 am and 11:00 pm. Each questionnaire was randomly delivered within this 16-hour time window, with a minimum of 30 minutes in between each notification.

### Multi-dimensional Experience Sampling

Participants received notifications on their phones through the Samply Research mobile application. All responses were made in reference to their thoughts, feelings, environment, location, and activities immediately before receiving the notification. This study used a 16-question mDES battery that has been used in prior studies [11, 13] (see Table 1). The mDES questions were always asked first and delivered in a random order. Participants then rated their stress on a 1-to-10 Likert scale. For the purpose of this study, responses to the stress question were not analyzed. Participants then answered questions about their physical and virtual social environment (Table 2). Finally, participants also indicated the type of physical location they were in (Table 3) and their primary activity (Table 4). The activity list was developed from the day reconstruction method [29] and modified based on the options in our prior studies [9, 30].

**Table 1.** mDES Questions.

| Dimension | Question | Scale Low | Scale High |
| --- | --- | --- | --- |
| Task | My thoughts were focused on an external task or activity: | Not at all | Completely |
| Future | My thoughts involved future events: | Not at all | Completely |
| Past | My thoughts involved past events: | Not at all | Completely |
| Self | My thoughts involved myself: | Not at all | Completely |
| People | My thoughts involved other people: | Not at all | Completely |
| Emotion | The emotion of my thoughts was: | Negative | Positive |
| Images | My thoughts were in the form of images: | Not at all | Completely |
| Words | My images were in the form of words: | Not at all | Completely |
| Sounds | My thoughts were in the form of sounds: | Not at all | Completely |
| Detailed | My thoughts were detailed and specific: | Not at all | Completely |
| Deliberate | My thoughts were: | Spontaneous | Deliberate |
| Problem | I was thinking about solutions to problems or goals: | Not at all | Completely |
| Intrusive | My thoughts were intrusive: | Not at all | Completely |
| Knowledge | My thoughts contained information I already knew (e.g., knowledge or memories): | Not at all | Completely |
| Absorption | I was absorbed in the contents of my thoughts: | Not at all | Completely |
| Distracting | My thoughts were distracting me from what I was doing: | Not at all | Completely |

*Note.* Participants rated statements on a 1-to-10 Likert scale.

**Table 2.** Social Environment Questions.

| Environment | Question | Environment Type |
| --- | --- | --- |
| Physical | Were you alone, or physically with other people? | Alone |
|  |  | Around people but not interacting with them |
|  |  | Around people and interacting with them |
|  | Where you alone, or physically with a pet? | Not with a pet |
|  |  | Around a pet but not interacting with them |
|  |  | Around a pet and interacting with them |
| Virtual | Were you alone, or virtually with other people? | Alone |
|  |  | Around people but not interacting with them (e.g., reading messages but not replying, being on a video call but not participating, etc.) |
|  |  | Around people and interacting with them (e.g., direct communication with another person by text, instant messaging, calling, or video calling, etc.) |

**Table 3.** Physical Environment Question.

| List Type | Question | Location List |
| --- | --- | --- |
| Location | Where were you? | Inside a home |
|  |  | Inside a shop |
|  |  | Inside a workplace |
|  |  | Inside (other) |
|  |  | Outside in a city or town |
|  |  | Outside in nature |
|  |  | Outside (other) |

*Note.* If participants selected “Inside (other),” or “Outside (other),” they were asked to specify their location.

**Table 4.** Primary Activity Question.

| List Type | Question | Activity List |
| --- | --- | --- |
| Activity | What were you doing? | Eating<br>Homework<br>Household chores<br>Listening to music<br>Napping or resting<br>Nothing or waiting<br>Personal exercise<br>Personal hygiene care<br>Physical leisure or sports<br>Reading<br>Shopping<br>Talking in person<br>Talking on the phone<br>Video-calling<br>Messaging by phone/device<br>Traveling or commuting<br>Using a computer or an electronic device<br>Walking the dog<br>Watching TV<br>Working<br>Other activity |

*Note.* If participants selected “Other activity,” they were asked to specify what they were doing. If participants selected “Using a computer or an electronic device,” they were asked to further specify their activity from a list of alternative activities that included “Social media: Passive scrolling,” “Social media: Active engagement,” “Gaming,” “Admin (e.g., banking, calendar, etc.),” and “Other.” If participants selected “Other,” they were asked to specify what they were doing).

## Analysis

### Data and Code Availability Statement

All custom code used to prepare data for analysis and figure development is available online at https://github.com/ThinCLabQueens and https://github.com/BridgMul10. Anonymized data has been uploaded to a publicly accessible database, Mendeley Data, and is available online at https://doi.org/10.17632/gfhwg3p8r6.1.

### Principal Component Analysis: mDES Questions

As in our prior studies (e.g. [8, 9, 12, 14, 30]), common patterns of thought (Figure 1) were identified by applying principal component analysis (PCA) using Varimax rotation to the data generated from responses to the 16 mDES questions (Table 1) using IBM SPSS Statistics (version 29). PCA was applied at the observation level and included 6776 observations. The loadings from the five components with an eigenvalue > 1 were retained for further analysis (see Table 5 and Figure 1).

**Figure 1.**
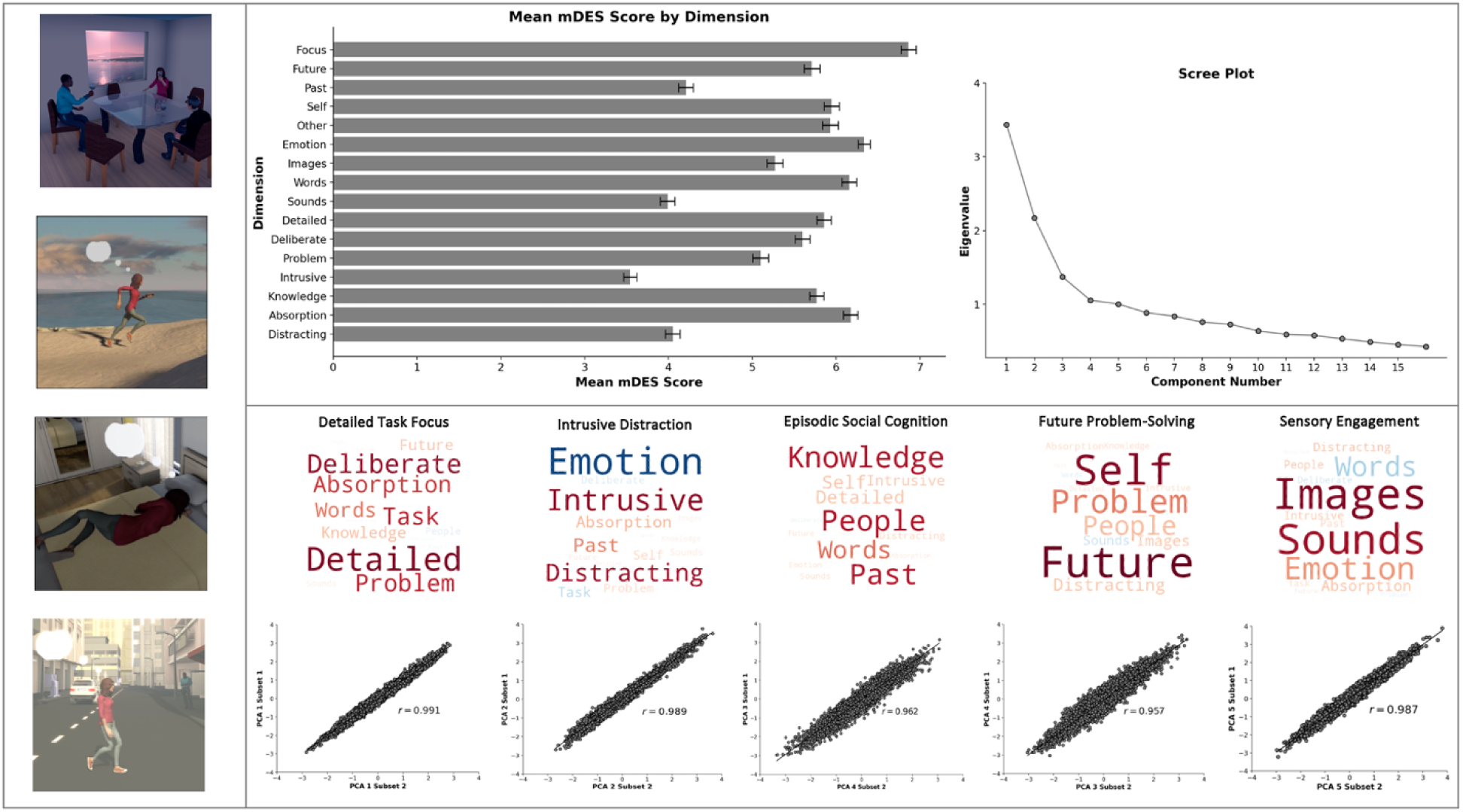
The Within Dataset Consistency of the Dimensional Structure of Ongoing Experience in Daily Life. *Note*. Upper horizontal panel: (Left) Bar graph describing the mean mDES score for each dimension. Error bars represent 99% confidence intervals (CIs). (Right) Scree plot generated from PCA of mDES questionnaire data. Lower horizontal panel: Word clouds describing the experiential features that contribute the most to the dimensions of experience as described by PCA (varimax rotation). Larger fonts describe features with more importance on the dimension (i.e., stronger loadings) and the font colour denotes direction of the loading (i.e., warm colours relate to positive loadings). For the purpose of exposition, these components are named based on their strongest features (from left to right): “Detailed Task Focus,” “Intrusive Distraction,” “Episodic Social,” “Future Problem-Solving,” and “Sensory Engagement.” Scatter plots below show the split half reliability of these dimensions (Y-axis: Subset 1; X-axis: Subset 2). Left vertical panel: Illustrations of different activities in daily life.

**Table 5.** PCA Loadings Generated from mDES Questions.

| Dimension | Detailed Task Focus | Intrusive Distraction | Episodic Social Cognition | Future Problem-Solving | Sensory Engagement |
| --- | --- | --- | --- | --- | --- |
| Task | .605 | -.207 | .017 | -.033 | .142 |
| Future | .228 | .068 | .106 | .797 | .064 |
| Past | -.021 | .408 | .636 | .054 | .169 |
| Self | .003 | .166 | .227 | .730 | -.009 |
| Person | -.079 | -.004 | .645 | .253 | .208 |
| Emotion | .018 | -.730 | .115 | .042 | .351 |
| Images | .017 | .048 | .032 | .216 | .742 |
| Words | .435 | .020 | .452 | -.092 | -.264 |
| Sounds | .051 | .080 | .127 | -.177 | .680 |
| Detailed | .748 | .038 | .224 | .102 | .034 |
| Deliberate | .702 | -.109 | -.059 | .009 | -0.137 |
| Problem | .609 | .146 | .010 | .412 | -0.052 |
| Intrusive | -.013 | .710 | .211 | .137 | 0.207 |
| Knowledge | .240 | .059 | .651 | .141 | -.002 |
| Absorption | .560 | .267 | .084 | .155 | .230 |
| Distracting | -.015 | .689 | .174 | .220 | .228 |

### Reliability Analysis: mDES Question Data

Component reliability analysis was conducted in IBM SPSS Statistics (version 29). mDES thought dimension data was randomly shuffled and divided into two halves (n = 3445 probes per sample). PCA (varimax) was applied to each subset separately and Pearson correlations were run to compare the component loadings generated from each subset with the overall solution. The higher the correlation between the two halves of the data to the overall sample, the more consistent the dimensional structure within the overall sample was.

### Linear Mixed Modeling: Generating “Thought-Activity” Mappings

To analyze contextual distributions of thought in relation to activities, we conducted a series of linear mixed models (LMMs), one with each thought component as the dependent variable and activity as the independent variable, to examine whether patterns of thought varied across activity categories. Restricted maximum likelihood (REML) was used as the estimation method and a variance components model was used as the covariance type. Participants were included as a random intercept. The parameter estimates for each activity in each model were saved for the eventual generation of activity word clouds that describe the experiential features that contribute to each dimension (Figure 2). This analysis is identical to that found in Mulholland et al. [9]. The parameter estimates were also saved to be used in reliability analysis (Figure 3).

**Figure 2.**
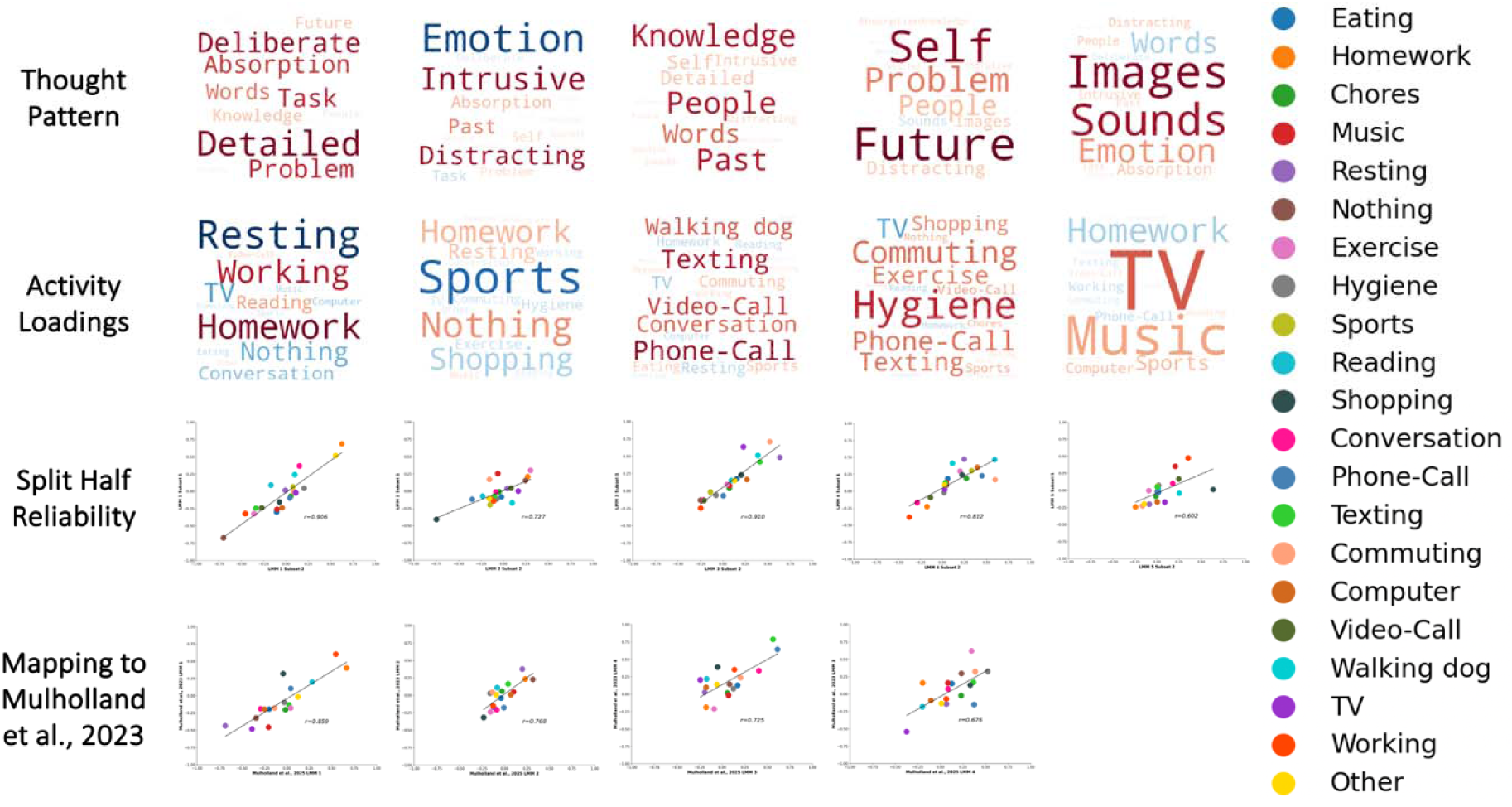
Consistency of Thought-Activity Mappings. *Note.* First row: “Thought Patterns.” Words represent PCA (varimax) loadings for mDES data and LMM loadings for primary activities. Larger fonts are features with more importance (i.e., higher loadings) and colour denotes features with a similar loading on the dimensions (i.e., warm colours relate to positive loadings). See Table 5-10 for specific component loadings. Second row: “Activity Loadings.” In these word clouds the large the font the stronger this activity loads on that thought component and activities with a similar colour load in the same direction. Third row: Scatter plots showing the within study consistency of the thought-activity mappings. (Y-axis: Activity loadings from Subset 1; X-axis: Activity Loadings from Subset 2). Fourth row: Scatter plots showing the between study consistency of the thought-activity mappings (Y-axis: Activity loadings from Mulholland et al. [7]; X-axis: Activity loadings in the current dataset).

**Figure 3.**
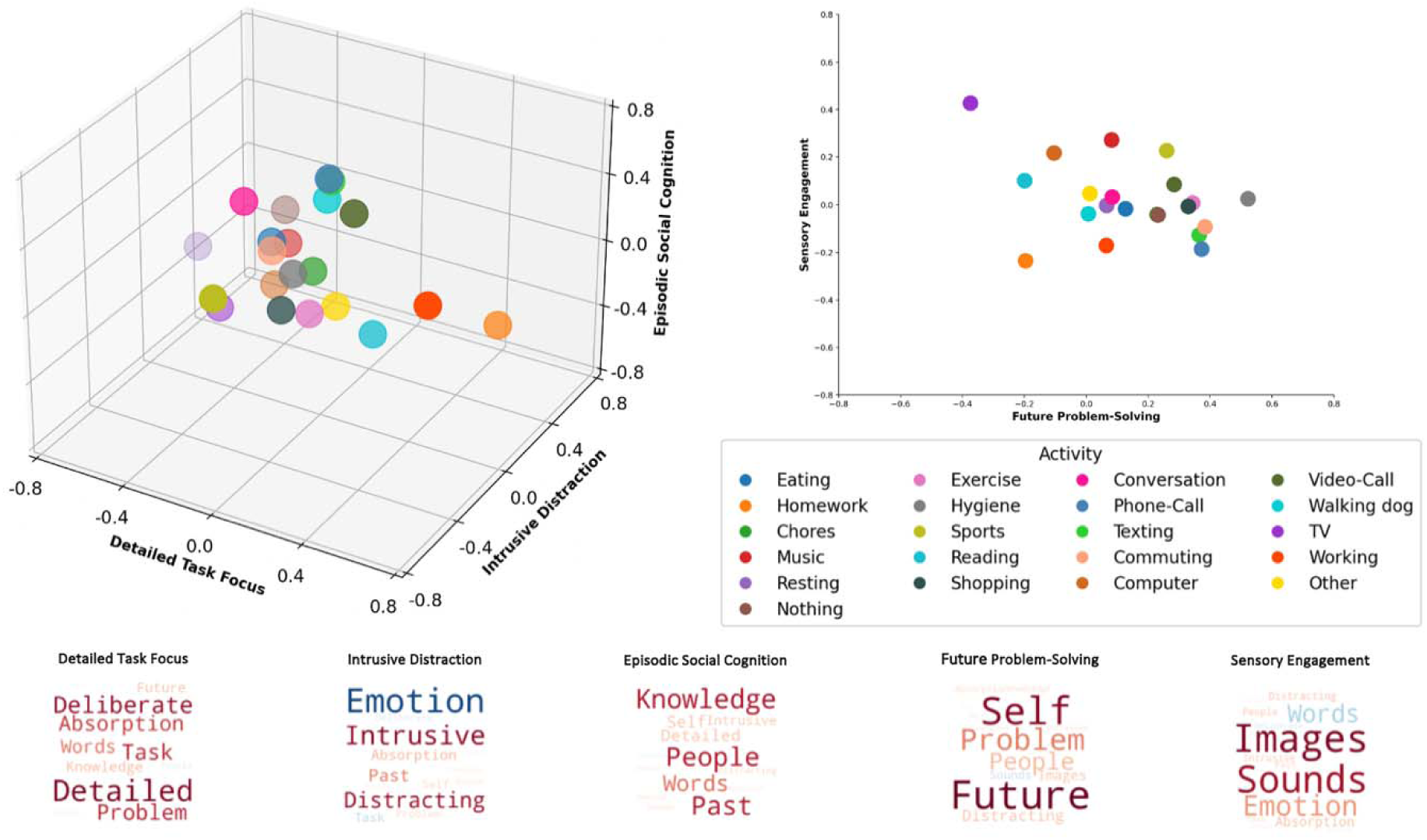
The Multivariate Landscape of Thought-Activity Mappings in Daily Life. *Note*. The thought-activity mapping revealed by our analysis are presented in two- and three-dimensional spaces to provide the opportunity to visualize the multivariate nature of the relationships. Left image: Thought-activity mappings in a three-dimensional “Thought Space.” Each point is an activity. X-axis is the activity loadings for “Detailed Task Focus”, the Y-axis is the activity loading on “Intrusive Distraction,” and the Z-axis are the activity loadings for “Episodic Social Cognition.” Right image: Thought-activity mappings in a two-dimensional thought space. The Y-axis is the activity loading on “Sensory Engagement” and the X-axis as the activity loadings on “Future Problem-Solving.” Lower panel: This shows the feature loadings for each dimension of the thought space in the form of word clouds. As described previously, larger fonts are activities with more importance (i.e., higher loadings on a dimension) and words in a similar colour denotes direction (i.e., warm colours relate to positive loadings).

**Table 6.** LMM Estimated Marginal Means for the “Detailed Task Focus” Component.

| Primary Activity | Mean | Std. Error | Lower Bound 95%<br>Confidence Interval | Upper Bound 95%<br>Confidence Interval |
| --- | --- | --- | --- | --- |
| Eating | -.198 | 0.044 | -.284 | -.112 |
| Homework | .664 | .033 | .599 | .729 |
| Chores | -.017 | .058 | -.130 | .096 |
| Music | -.205 | .062 | -.326 | -.083 |
| Resting | -.686 | .043 | -.770 | -.602 |
| Nothing | -.344 | .057 | -.455 | -.233 |
| Exercise | .042 | .080 | -.114 | .199 |
| Hygiene | -.029 | .060 | -.146 | .088 |
| Sports | -.149 | .127 | -.398 | .099 |
| Reading | .281 | .080 | .124 | .438 |
| Shopping | -.042 | .094 | -.226 | .143 |
| Conversation | -.293 | .043 | -.377 | -.209 |
| Phone-call | .043 | .102 | -.157 | .243 |
| Video-call | .168 | .089 | -.007 | .344 |
| Texting | .025 | .084 | -.140 | .189 |
| Commuting | -.139 | .077 | -.291 | .012 |
| Computer | -.250 | .042 | -.333 | -.167 |
| Walking dog | -.005 | .165 | -.329 | .318 |
| TV | -.388 | .042 | -.470 | -.306 |
| Working | .544 | .052 | .441 | .646 |
| Other | .122 | .043 | .038 | .206 |

**Table 7.**
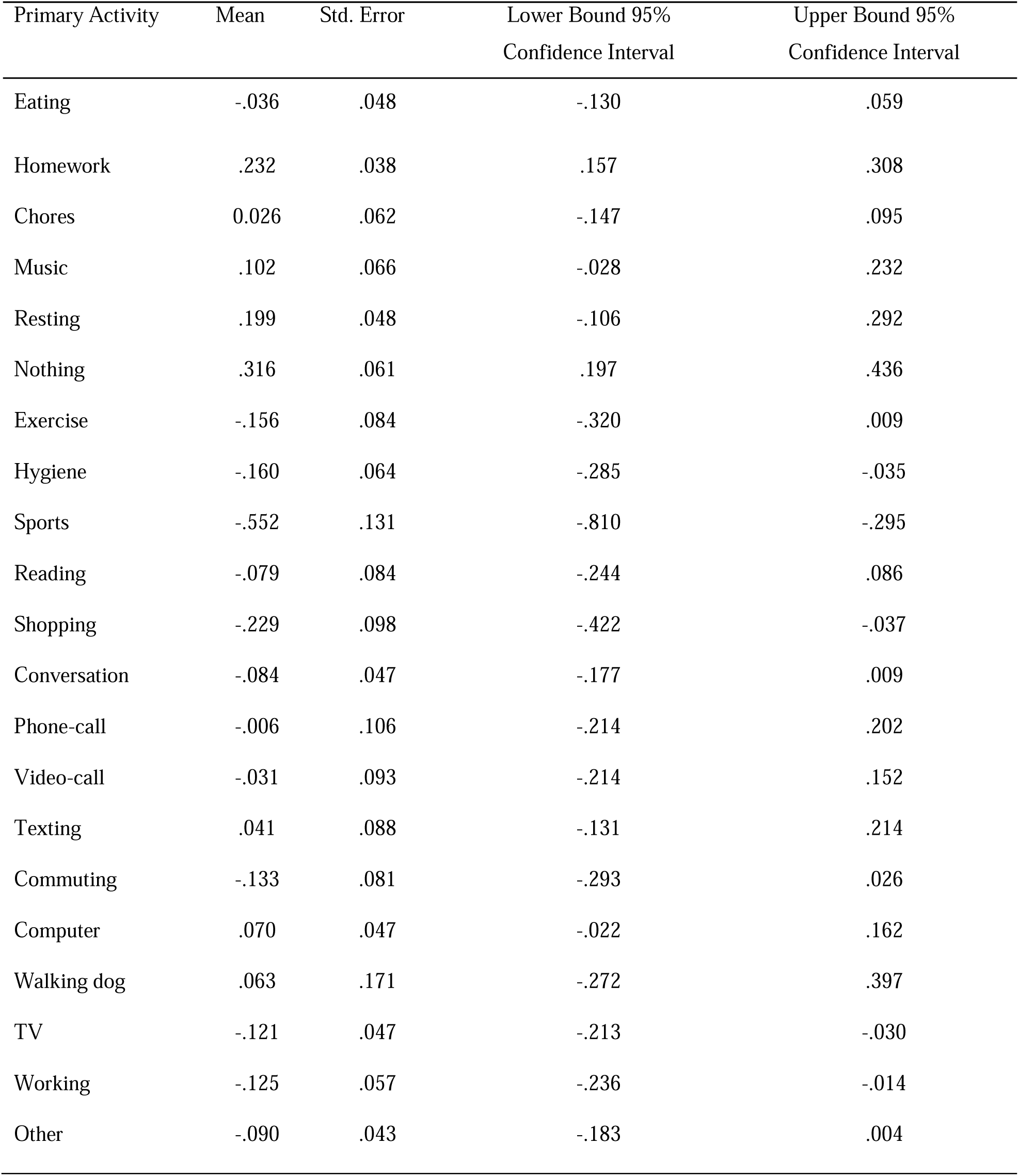
LMM Estimated Marginal Means for the “Intrusive Distraction” Component.

**Table 8.** LMM Estimated Marginal Means for the “Episodic Social Cognition” Component.

| Primary Activity | Mean | Std. Error | Lower Bound 95%<br>Confidence Interval | Upper Bound 95%<br>Confidence Interval |
| --- | --- | --- | --- | --- |
| Eating | 0.170 | 0.047 | 0.078 | 0.262 |
| Homework | -0.178 | 0.036 | -0.247 | -0.108 |
| Chores | 0.057 | 0.062 | -0.063 | 0.178 |
| Music | 0.070 | 0.066 | -0.060 | 0.200 |
| Resting | -0.194 | 0.046 | -0.284 | -0.104 |
| Nothing | 0.082 | 0.061 | -0.036 | 0.201 |
| Exercise | -0.089 | 0.085 | -0.256 | 0.078 |
| Hygiene | 0.124 | 0.064 | -0.001 | 0.249 |
| Sports | 0.195 | 0.135 | -0.070 | 0.459 |
| Reading | -0.171 | 0.085 | -0.338 | -0.003 |
| Shopping | -0.050 | 0.100 | -0.246 | 0.147 |
| Conversation | 0.405 | 0.046 | 0.315 | 0.494 |
| Phone-call | 0.610 | 0.109 | 0.397 | 0.823 |
| Video-call | 0.469 | 0.095 | 0.283 | 0.656 |
| Texting | 0.561 | 0.089 | 0.385 | 0.736 |
| Commuting | 0.202 | 0.083 | 0.040 | 0.364 |
| Computer | -0.178 | 0.045 | -0.267 | -0.089 |
| Walking dog | 0.429 | 0.176 | 0.085 | 0.773 |
| TV | -0.243 | 0.045 | -0.331 | -0.155 |
| Working | 0.136 | 0.056 | 0.026 | 0.245 |
| Other | -0.058 | 0.046 | -0.148 | 0.032 |

**Table 9.**
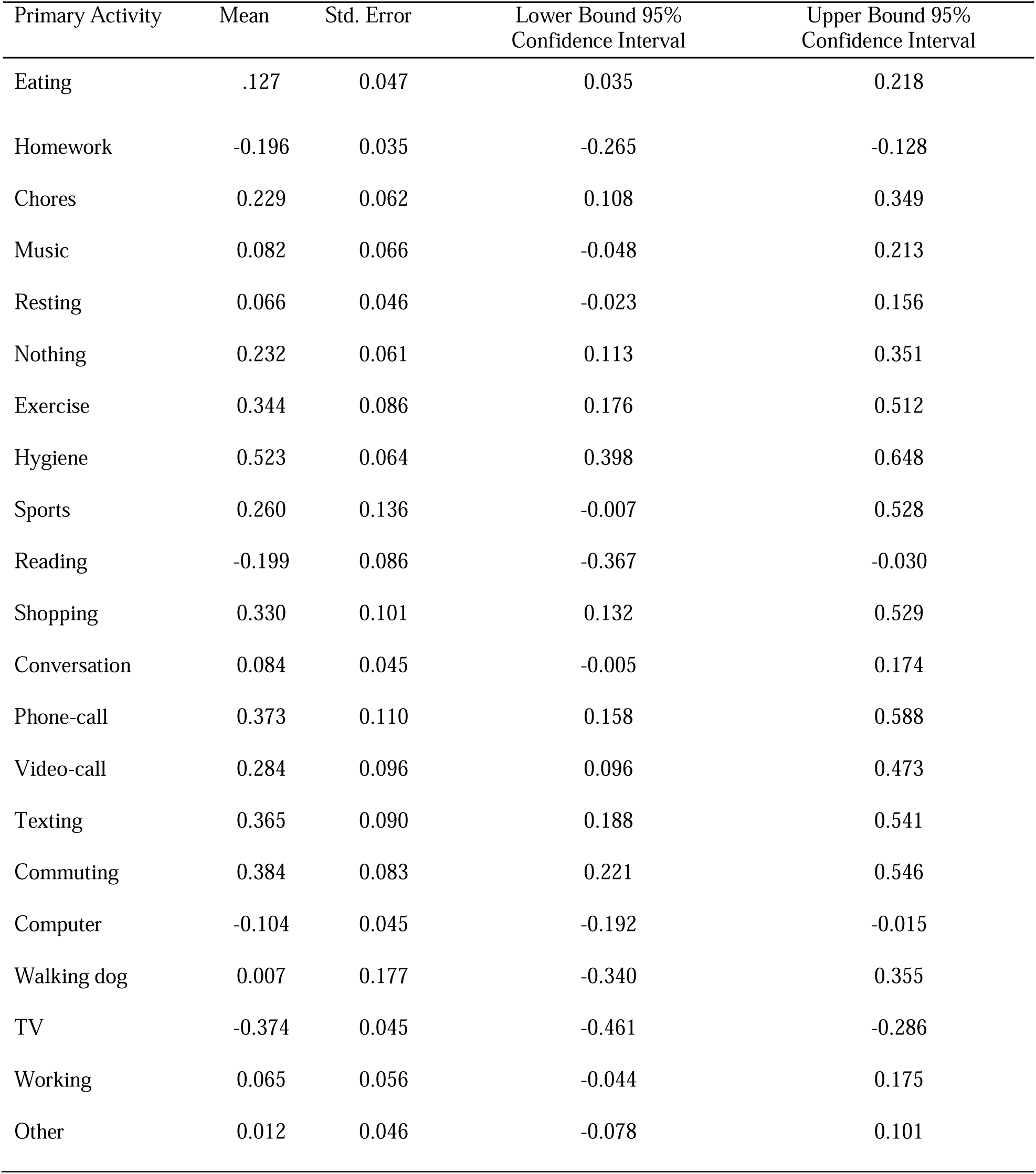
LMM Estimated Marginal Means for the “Future Problem-Solving” Component.

| Primary Activity | Mean | Std. Error | Lower Bound 95%<br>Confidence Interval | Upper Bound 95%<br>Confidence Interval |
| --- | --- | --- | --- | --- |
| Eating | .127 | 0.047 | 0.035 | 0.218 |
| Homework | -0.196 | 0.035 | -0.265 | -0.128 |
| Chores | 0.229 | 0.062 | 0.108 | 0.349 |
| Music | 0.082 | 0.066 | -0.048 | 0.213 |
| Resting | 0.066 | 0.046 | -0.023 | 0.156 |
| Nothing | 0.232 | 0.061 | 0.113 | 0.351 |
| Exercise | 0.344 | 0.086 | 0.176 | 0.512 |
| Hygiene | 0.523 | 0.064 | 0.398 | 0.648 |
| Sports | 0.260 | 0.136 | -0.007 | 0.528 |
| Reading | -0.199 | 0.086 | -0.367 | -0.030 |
| Shopping | 0.330 | 0.101 | 0.132 | 0.529 |
| Conversation | 0.084 | 0.045 | -0.005 | 0.174 |
| Phone-call | 0.373 | 0.110 | 0.158 | 0.588 |
| Video-call | 0.284 | 0.096 | 0.096 | 0.473 |
| Texting | 0.365 | 0.090 | 0.188 | 0.541 |
| Commuting | 0.384 | 0.083 | 0.221 | 0.546 |
| Computer | -0.104 | 0.045 | -0.192 | -0.015 |
| Walking dog | 0.007 | 0.177 | -0.340 | 0.355 |
| TV | -0.374 | 0.045 | -0.461 | -0.286 |
| Working | 0.065 | 0.056 | -0.044 | 0.175 |
| Other | 0.012 | 0.046 | -0.078 | 0.101 |

**Table 10.** LMM Estimated Marginal Means for the “Sensory Engagement” Component.

| Primary Activity | Mean | Std. Error | Lower Bound 95%<br>Confidence Interval | Upper Bound 95%<br>Confidence Interval |
| --- | --- | --- | --- | --- |
| Eating | -0.017 | 0.049 | -0.113 | 0.080 |
| Homework | -0.235 | 0.041 | -0.315 | -0.155 |
| Chores | -0.041 | 0.061 | -0.160 | 0.079 |
| Music | 0.272 | 0.065 | 0.145 | 0.400 |
| Resting | -0.003 | 0.049 | -0.098 | 0.092 |
| Nothing | -0.043 | 0.060 | -0.161 | 0.075 |
| Exercise | 0.007 | 0.081 | -0.152 | 0.165 |
| Hygiene | 0.025 | 0.063 | -0.098 | 0.148 |
| Sports | 0.227 | 0.125 | -0.018 | 0.471 |
| Reading | 0.101 | 0.081 | -0.059 | 0.260 |
| Shopping | -0.007 | 0.094 | -0.191 | 0.177 |
| Conversation | 0.032 | 0.048 | -0.063 | 0.126 |
| Phone-call | -0.186 | 0.101 | -0.385 | 0.013 |
| Video-call | 0.085 | 0.090 | -0.091 | 0.261 |
| Texting | -0.127 | 0.085 | -0.293 | 0.039 |
| Commuting | -0.093 | 0.079 | -0.248 | 0.061 |
| Computer | 0.217 | 0.048 | 0.123 | 0.311 |
| Walking dog | -0.038 | 0.162 | -0.355 | 0.279 |
| TV | 0.427 | 0.048 | 0.333 | 0.520 |
| Working | -0.171 | 0.057 | -0.282 | -0.060 |
| Other | 0.047 | 0.049 | -0.048 | 0.143 |

### Reliability Analysis: Within Data Set Consistency of Thought-Activity Mappings

Primary activity data reliability analysis was conducted in IBM SPSS Statistics (version 29). As before, component scores were split into two subsets (n = 3445 probes per subset). We conducted a series of LMMs, with thought components for each subset as the dependent variable and activity as the independent variable. Restricted maximum likelihood (REML) was used as the estimation method and a variance component model was used as the covariance type.

Participants were included as a random intercept. Pearson correlations were run on the parameter estimates generated from each subset for each activity. This analysis allowed us to estimate whether the results of our whole sample LMM are generalizable to sub-samples of the data.

### Reliability Analysis: Comparing Thought-Activity Mappings Across Studies

Reliability analysis was conducted in IBM SPSS Statistics (version 29). The LMM parameter estimates from Mulholland et al. [31] contained a sample of n = 1451 probes, and the LMM parameter estimates from our current study contained a sample of n = 6889 probes.

Pearson correlations were run on the parameter estimates for overlapping activity from each dataset. This analysis allowed us to estimate whether the results of our whole sample LMM in the current data are generalizable to those seen in the prior dataset.

### Linear Mixed Modeling: Social Environment Data

To analyze contextual distributions of thought in relation to socialization, we conducted a series of LMMs, one with each thought component as the dependent variable and social environment as the independent variable (i.e., physical environment with people, physical environment with a pet, or virtual environment with people were all options as independent variables). These analyses allowed us to examine whether patterns of thought varied meaningfully across social environments. Restricted maximum likelihood (REML) was used as the estimation method, and a variance components model was used as the covariance type. Participants were included as a random intercept.

### Multiple Regression: Linking mDES to Traits

To analyze whether mDES questions can differentiate individuals based on underlying traits, we performed a series of multiple regressions with the average PCA component scores for each individual in the overall “Thought Space” as the dependent variable and trait questionnaires as explanatory variables. Trait questionnaires included in this analysis include the AQ, ASRS, MADRS, OASIS, and WHOQOL-BREF, all of which were z-scored. The WHOQOL-BREF questionnaire was subdivided into four categories based on scoring domains: physical health, psychological health, social relationships, and environment. mDES questionnaire data was reduced to a single score for each participant.

## Results

### Patterns of Ongoing Thought

First, mean scores for each thought dimension measured were calculated and are shown in the bar plots in Figure 1. Next, as is standard in our laboratory work (e.g. [8, 12, 14, 30]) the mDES data was decomposed using PCA (varimax rotation) to reveal the dimensions that best described the patterns of ongoing thought reported by the participants. Based on eigenvalue > 1, five components were selected for further analysis (see Figure 1 for scree plot). PCA loadings (Table 5) from the five components were used to generate thought word clouds (Figure 1).

Components were named based on mDES thought dimensions that dominated their composition. Component 1 (21.5% of variance) was labelled “Detailed Task Focus” because loadings were high for “detailed,” “deliberate,” “problem,” and “task” (Figure 1). Component 2 (13.6% of variance) was labelled “Intrusive Distraction” because loadings were high for “(negative) emotion,” “intrusive,” and “distracting” (Figure 1). Component 3 (8.6% of variance) was labelled “Episodic Social Cognition” because loadings were high for dimensions such as “knowledge,” “person,” and “past” (Figure 1). Component 4 (6.6% of variance) was labelled “Future Problem-Solving” because loadings were high for dimensions such as “future,” “self,” and “problem” [as in problem solving] (Figure 1). Component 5 (6.3% of variance) was labelled “Sensory Engagement” because loadings were high for dimensions such as “images,” and “sounds” (Figure 1). Please note that these terms are used for convenience to summarize the features that characterize each component. They do not constitute the only labels which could be applied to these patterns.

### Component Reliability

To understand the robustness of the dimensions within our sample, we conducted a split-half reliability analysis. In this analysis, the data was divided into two random samples (n = 3445 probes per sample) and then examined to determine how the components generated in each half of the data related to each other. We used the robustness of the solutions across PCAs with 3-, 4-, and 5-component solutions as a complementary method to determine the best solution for the entire sample. The mean correlation for the set of the homologous pairs from each solution was calculated with a higher score reflecting the most reproducible components. The 3-, 4-, and 5-component solutions all produced highly reliable results, with average homologue similarities scores of *r* = .996 (3-component solution, range *r* = .994-.995, Figure 1), *r* = .93525 (4-component solution, range *r* = .869-.978, Figure 1), and *r* = 0.9772 (5-component solution, range *r* =.957-.991, Figure 1). Since the similarity was high for 3-, 4-, and 5-component solutions, we focused our analysis on the 5-component solution.

### Thought-Activity Mappings

Having identified the stability of the dimensions identified within this dataset, we examined if mDES provides a reliable way to map cognition in daily life onto activities that are being performed. To this end, we ran a series of LMMs in which the activities are the explanatory variables and the mDES scores for each dimension from the common “Thought-Space” are the dependent variables. In each case we found a significant association between reported patterns of thought and activities (“Detailed Task Focus” (*F*(21, 6639.50) = 74.08, *p* <.001); “Intrusive Distraction” (*F*(21, 6591.47) = 10.21, *p* <.001); “Episodic Social Cognition” (*F*(21, 6632.08) = 19.08, *p* <.001); “Future Problem-Solving” (*F*(21, 6648.67) = 18.26, *p* <.001); “Sensory Engagement” (*F*(21, 6559.04) = 16.82, *p* <.001)). To visualize these thought-activity mappings we generated a set of word clouds based on the estimated marginal means for reported activities in each component (Figure 2). “Detailed Task Focus” was high when doing homework and working (paid or volunteer) and lowest when napping or resting, watching TV, and doing nothing or waiting. “Intrusive Distraction” was highest when doing nothing or waiting, napping or resting, and doing homework, and lowest during physical leisure or sports, and shopping. “Episodic Social” features were highest when talking on the phone, messaging by phone/device, video-calling, walking the dog, talking in person, traveling or commuting, and physical leisure or sports, and lowest when watching TV, napping or resting, using a computer or electronic device, and doing homework. “Future Problem-Solving” was higher during hygiene activities, texting, commuting, phone-call, exercise, and shopping, and lowest when watching TV, reading, and doing homework. Lastly, “Sensory Engagement” had high loadings when watching TV, listening to music, playing sports, and using a computer, and low loadings when doing homework, talking on the phone, and working.

Next, we investigated the robustness of the thought-activity mappings produced by mDES. Our first analysis examined the consistency of their mappings within the current data by performing a split-half reliability analysis with current data. We created two subsets of our data (n = 3445 probes per subset). In each case, we found a significant association between each of the reported thought patterns and activities in each subset (“Detailed Task Focus” subset 1 (*F*(21, 3284.10) = 36.96, *p* <.001) and subset 2 (*F*(21, 3333.340) = 36.101, *p* <.001); “Intrusive Distraction” subset 1 (*F*(21, 3274.05) = 4.63, *p* <.001) and subset 2 (*F*(21, 3285.57) = 6.53, *p* <.001); “Episodic Social Cognition” subset 1 (*F*(21, 3323.14) = 8.92, *p* <.001) and subset 2 (*F*(21, 3271.06) = 10.32, *p* <.001); “Future Problem-Solving” (*F*(21, 3285.56) = 10.02, *p* <.001) and subset 2 (*F*(21, 3294.73) = 9.06, *p* <.001); “Sensory Engagement” subset 1 (*F*(21, 3242.37) = 9.69, *p* <.001) and subset 2 (*F*(21, 3239.06) = 8.13, *p* <.001)).

To understand whether the thought-activity mappings in daily life were similar for each subset of the data the loadings of each activity on each component each component was correlated (see Methods). Correlations between the LMM activity estimated marginal means in each subset ranged from *r* = .60-.91, with the highest reproducibility for the “Detail Task Focus” component and the lowest for the “Sensory Engagement” component. This shows a reasonably high degree of consistency in the feature loadings for each dimension of experience within the current sample.

Next, we examined how the thought-activity mappings from the current study map onto those from the homologous components from Mulholland et al. [9]. To this end, we used Pearson Correlation to compare the activity mappings from our prior study to the mappings found in our omnibus sample. Mulholland et al. [9] did not include a “Sensory Engagement” component (because the modality questions did not include sounds), so only four components were compared. This analysis identified correlations with a range of *r* = .67-.86, with the strongest mappings for “Detailed Task Focus” and the lowest for “Future Problem-Solving.” This analysis shows a reasonably high degree of overlap between the thought activity mappings as described in these data and those seen in our prior study [9].

One feature of our analysis is that it allows us to understand experience within each activity as a linear combination of different thought components which can be represented graphically by an activities location in a multivariate “Thought Space.” To visualize the data this way, the estimated marginal means for each activity derived from the LMMs and plotted against each mDES components for the common “Thought-Space.” We present these data a three-dimensional space constructed using the three most reliable components (i.e., “Detailed Task Focus,” “Intrusive Distraction,” and “Episodic Social Cognition”) as the dimensions. We also present a two-dimensional space that describes the relationship between activities in terms of their weighting for “Future Problem-Solving” and “Sensory Engagement.” These are both presented in (, which show how certain activities occupy extreme values on multiple components. For example, “homework” is high on both the “Detailed Task Focus” and “Intrusive Distraction” components but low on “Episodic Social Cognition,” “Future Problem-Solving,” and “Sensory Engagement.”

### Thought Dimensions and Social Environments

Another way to examine the robustness of mDES as a tool for mapping cognition in daily life by measuring the similarity in how the thought patterns map onto social environments in daily life. Prior studies (e.g. [9]) have found that being alone compared to being with other people have important implications for a persons’ thought patterns, in particular facilitating patterns of thought with social and episodic features. We ran a series of LMMs in which different social environments (Table 2) were the explanatory variable and the mDES scores from the common “Thought Space” the dependent variables. We found a significant association between reported patterns of thought and the physical social environments for each component. “Detailed Task Focus” (*F*(3, 6721.87) = 16.06, *p* <.001) was lowest when with people and interacting with them (*M* = -.08, 95% CI [-.08, -.13]) and highest when around people but not interacting with them (*M* = .14, 95% CI [.08,-.21]). “Intrusive Distraction” (*F*(3, 6656.39) = 37.54, *p* <.001) was lowest when with people and interacting (*M* = -.15, 95% CI [-.22, -.08]) and highest when alone (*M* = .11, 95% CI [.05,.18]). “Episodic Social Cognition” (*F*(3, 6699.28) = 56.29, *p* <.001) was highest when around people and interacting (*M* = .22, 95% CI [.16,.28]) and lowest when around people but not interacting (*M* = -.15, 95% CI [-.08,-.22]). “Future Problem-Solving” (*F*(3, 6716.79) = 2.79, *p* = .039) was lowest when around people and interacting with them (*M* = −0.46, 95% CI [-.11,.02]) and highest when around people but not interacting (*M* = .04, 95% CI [-.02,.15]). Finally, “Sensory Engagement” (*F*(3, 6619.47) = 5.30, *p* = .001) was highest when alone (*M* = .03, 95% CI [-.08,.02]). A broadly similar pattern was found for our analysis of the consequence of virtual environments. “Detailed Task Focus” (*F*(3, 6705.72) = 2.67, *p* = .046) was higher when around people and not interacting (*M* = .05, 95% CI [-.03,.14]) and lowest when alone (*M* = -.03, 95% CI [-.83,.02]). “Intrusive Distraction” (*F*(3, 6695.70) = 11.53, *p* <.001) was least prevalent when participants were around people and interacting virtually (*M* = -.11, 95% CI [-.19,-.03]) and highest when around people but not interacting virtually (*M* = .07, 95% CI [-.02, .16]). “Episodic Social Cognition” (*F*(3, 6715.54) = 39.37, *p* <.001) was highest when around people and interacting virtually (*M* = .27, 95% CI [.19,.34]) and lowest when alone (*M* = -.07, 95% CI [.19,.33]). “Future Problem-Solving” (*F*(3, 6709.60) 9.55, *p* <.001) was highest when around people but not interacting (*M* = .15, 95% CI [.07,.22]) and lowest when around people but not interacting virtually with another person (*M* = -.03, 95% CI [-.12,.05]). Finally, “Sensory Engagement” (*F*(3, 6663.16) = 2.89, *p* = .03) was highest when alone (*M* = .01, 95% CI [-.05,.08]) and lowest around people but not interacting virtually (*M* = -.003, 95% CI [-.09,.09]). These analyses replicate prior observations that found a mapping of social cognition onto social situations (e.g. [9, 30]) but extends this to establish that such situations also reduce intrusive features of cognition.

### Thought Dimensions and Traits

Having established a high level of consistency between the thought-activity mappings seen in the current study and prior studies [9, 30], the final goal of our study was to investigate whether mDES dimensions can differentiate people based on traits associated with mental health and well-being. To do so, we performed a series of multiple regressions with the average loading of each individual on each dimension of the common “Thought Space” as the dependent variables and their score on the trait questionnaires as explanatory variables. Results indicated the location of each individual on the “Intrusive Distraction” dimension (*F*(8, 252) = 8.35, *p* = <.001; see Table 11 for results for each component) could be predicted based on high levels of anxiety (Beta = 0.23, *p* = .003) and low levels of social well-being (Beta = −0.17, *p* =.010). This analysis, therefore, shows a clear mapping between patterns of “Intrusive Distraction” with higher levels of anxiety and lower social well-being (Figure 4).

**Figure 4.**
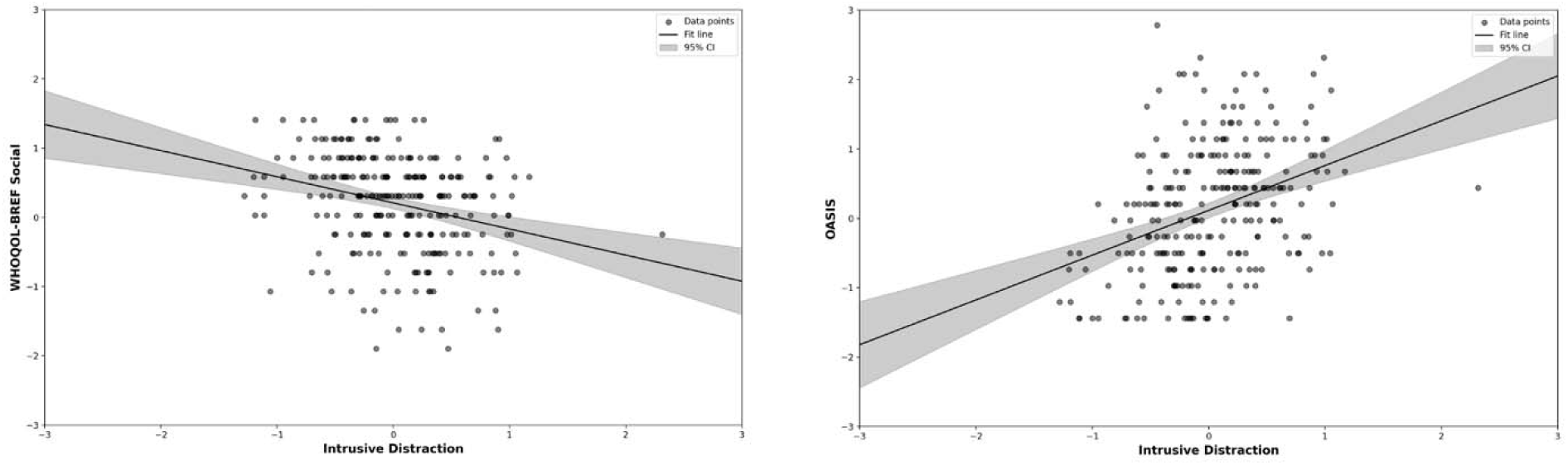
mDES Questionnaire Data and Trait Data Multiple Regression Results. *Note*. Left panel: Scatterplot with WHOQOL-BREF social relationships (z-scored) on the y-axis and “Intrusive Distraction” PCA component scores on the x-axis. Right panel: Scatterplot with OASIS (z-scored) on the y-axis and “Intrusive Distraction” PCA component scores on the x-axis. For both scatterplots, all values were mean z-scored per participant.

**Table 11.** Mental Health and Well-Being Multiple Regression Results Per PCA Component Scores.

| Component | Model | Sum of Squares | df | Mean Square | F | Sig. |
| --- | --- | --- | --- | --- | --- | --- |
| Detailed Task | Regression | .660 | 8 | .083 | .444 | .894 |
| Focus | Residual | 46.866 | 252 | .186 |  |  |
|  | Total | 47.526 | 260 |  |  |  |
| Intrusive | Regression | 15.071 | 8 | 1.884 | 8.348 | <.001 |
| Distraction | Residual | 56.869 | 252 | .226 |  |  |
|  | Total | 71.940 | 260 |  |  |  |
| Episodic Social | Regression | 2.127 | 8 | .266 | 1.085 | .374 |
| Cognition | Residual | 61.773 | 252 | .245 |  |  |
|  | Total | 63.900 | 260 |  |  |  |
| Future | Regression | 3.129 | 8 | .391 | 1.806 | .076 |
| Problem- | Residual | 54.569 | 252 | .217 |  |  |
| Solving | Total | 57.698 | 260 |  |  |  |
| Sensory | Regression | 1.248 | 8 | .156 | .415 | .911 |
| Engagement | Residual | 94.767 | 252 | .376 |  |  |
|  | Total | 96.015 | 260 |  |  |  |

## Discussion

Our study sought to investigate the underlying relationships between patterns of reported thought, behavior, and measures of mental health in daily life. First, we investigated the stability of the patterns of thoughts reported in our study, establishing a reasonable range of consistency for a five-dimensional solution. These dimensions were “Detailed Task Focus,” “Intrusive Distraction,” “Episodic Social,” “Future Problem-solving,” and “Sensory Engagement.” Second, we examined the reproducibility of how activities in daily life map onto these dimensions. The thought-activity mappings were both consistent within these data and were similar to the thought patterns described by Mulholland et al. [9] (Figure 2), who administered mDES in daily life to a different sample of participants. We also replicated patterns in prior studies that show, for example, that patterns of “Episodic Social Cognition” are higher when individuals are interacting either physically or virtually observed in both prior studies [9], a pattern of thought that was also reduced during COVID lockdowns in the United Kingdom because of reductions in the opportunities for socialization [30]. Taken together these analyses show that the “Thought Spaces” generated by the application of mDES to daily life provide a reproducible measure ongoing thought and that these organize the activities we engage in as we go about our daily lives in a meaningful manner (see Figure 3). It is important to note that populations in both studies are undergraduate students, so this reproducibility highlights that if mDES is employed within a similar group of individuals, then it captures thought patterns that are reasonably consistent in their profile and their associations with activities. Further studies are needed to understand how thought-activity mappings vary across cultures. However, it is possible that the observed thought patterns, as well as their links to activities or social contexts, could conceivably change if they were measured in different cultures or age groups.

### Links Between Thought Patterns in Daily Life and Psychological Well-Being

The main goal of the study was to understand the links between patterns of thought in daily life and mental health. We found that patterns of thought that loaded heavily on “Intrusive Distraction” were associated with greater levels of anxiety and less social support (Figure 4). Importantly this thought pattern showed a reasonable degree of consistency in terms of how it mapped onto activities within this sample (*r* = .77) and with our prior study (*r* = .73) highlighting that this thought component has a reasonably consistent thought-activity mapping.

Contemporary perspectives that emphasize patterns of thought often have an important social component (e.g., [32]). One important contribution of our study is that our analysis suggests that higher levels of “Intrusive Distraction” are also linked to social processes. As well as linking intrusive thought to the relatively low social well-being, our analysis also shows that this pattern of thought tends to dominate solo activities (such as exercise, doing nothing, and homework) and is relatively absent during social interactions, either virtually or in person.

Interestingly, despite intrusive thought being relatively high during exercise, this component was relatively low during sports, providing further evidence that social contexts may reduce levels of unpleasant intrusive experiences. Importantly, these data are consistent with prior studies that shown links between anxiety and increased levels of mind-wandering (e.g., [33]), a self-generated state that is likely to have distracting features [15, 34]. Altogether, therefore, these data suggest that unpleasant intrusive thoughts may be promoted by loneliness and social isolation, a situation that may promote personal worries, often known as current concerns [35]. It is possible based on our study that interventions that increase opportunities for socialization may be an important way to help individuals who suffer from anxiety.

### Open Questions

As well as detailing links between thinking in daily life and well-being, our study also raises a number of important open questions. For example, the consistency of the thought-activity mappings suggests that in any given population there may be systematic relationships between what people think and what they do in daily life. Based on these results, it may be possible to develop an understanding of the activities in daily life that are most related to different features of a person’s thought content, a taxonomy of mappings between thoughts and activities which, as a discipline, we currently lack. It is worth noting that the capacity for mDES to map thoughts in the lab and in daily life [36] makes it a useful tool for this goal. The relative ease with which we can administer mDES in daily life makes it possible to study a relatively large and socio-culturally diverse population (e.g. older adults or individuals from a different culture). Thus while our study shows that mDES is reliable, the specific results presented in this study may not generalize to different cultures, ethnicities, or ages, our methodology of smartphone sampling is free and easy to use, so future studies can address this gap by broadening the scope of their sampling population in a reasonably accessible manner. It is also important to note that there were a number of instances of incorrect and/or missing demographic data in our data which should be considered a limitation of our current study.

It is also likely that the battery of questions we used in our study could be improved. In our study, we used a combination of mDES questions that had been used previously in both lab-based and daily-life-based settings, as well as new mDES questions based on previously identified limitations. Mulholland et al. [9] noted that including a single modality question does not allow participants to describe their conscious experience of an activity fully. In that study, a component that was common while listening to music was dominated by thoughts with images as a key feature (in that mDES battery there were no questions about sounds). In light of this discovery, we decided to replace the modality probe with three questions relating to an individual’s sensory experience (i.e., images, words, and sounds). The inclusion of these three questions most likely led to the discovery of the “Sensory Engagement” component in the current study. It is important to note that a similar component was observed in the context of movie watching (using the same set of items as was applied in the current study [11]) and was associated with periods of films when brain activity was activated within sensory systems (both auditory and visual). This demonstrates that it is possible to optimize the questions included in mDES approaches that could, in the future, provide more accurate ways for participants to describe their conscious experience. In the same vein, there are likely to be ways to improve the measures of mental health we employed in this analysis. The health and wellness surveys were selected based on a widespread, general approach to health and wellness, rather than clinical diagnosis. In the future, more accurate or informative questionnaires could be selected that would provide a better understanding of how our thoughts in daily life impact upon our mental health.

## Acknowledgements

This research is supported by (i) a National Science and Engineering Research Council (NSERC) graduate fellowship awarded to Bridget Mulholland, (ii) an award from the Government of Canada’s New Frontiers in Research Fund (NFRF) (grant ID: NFRF-2021-00183) to Dr. Jonathan Smallwood and Dr. Jeffrey D. Wammes, and (iii) an NSERC Discovery Grant awarded to Dr Jonathan Smallwood (Grant ID: 2023-03496).

